# Prior experience dynamically determines differential cue weighting in the medial entorhinal cortex

**DOI:** 10.64898/2026.07.22.739574

**Authors:** Pascal Klein, Beate Throm, Kevin Allen, Hannah Monyer

## Abstract

Spatial navigation depends on integrating multiple cue sets whose influence is determined primarily by their inferred reliability. Here we investigated how cue weighting in the medial entorhinal cortex is updated based on previous experience at different time scales (minutes vs. days). In mice recorded in light and darkness while manipulating coherence between distal room cues and local arena cues, grid cells displayed stable firing fields in darkness only when prior light experience maintained coherent cue relationships. Grid activity remained anchored to local cues, and stability continuously increased by repeated exposure. In contrast, mice that experienced cue mismatch in light showed markedly reduced grid stability in darkness. Strikingly, re-establishing cue coherence produced only partial recovery, whereas a single exposure to cue mismatch abolished reliance on local cues. The reference frame shift at different time scales reflects a dynamic balance between flexibility and stability determined by a conservative cue weighting strategy.

## Introduction

Navigation in complex and dynamically changing environments requires the integration of external sensory cues (e.g. visual landmarks) with self-motion signals. The process by which animals rely on self-motion cues (such as vestibular input, proprioception, and copies of the motor efference) for navigation is referred to as path integration^1^. Because path integration is inherently susceptible to error accumulation in the absence of external reference points, sensory landmarks can provide critical error-correction signals^2–4^. However, sensory cues do not invariably dominate navigation behavior. Rather, the relative influence of self-motion and sensory cues depends on their reliability and behavioral context^5,6^. Likewise, when different sensory cues conflict, animals dynamically weight individual cues according to their salience and stability, a process conserved across species including insects, birds, rodents, non-human primates, and humans^7–15^.

At the neural level, differential cue weighting is reflected in distinct representations across the parahippocampal formation and associated brain regions. Head direction (HD) cells in the anterodorsal thalamic nucleus (ADN) are strongly anchored to distal landmarks, yet they can maintain directional tuning in the absence of visual input, presumably through self-motion signals^16–18^. As the HD system is thought to operate through ring-attractor dynamics^19–22^, HD cells typically rotate coherently with a dominant cue set. In contrast, hippocampal place cells can exhibit heterogeneous responses under cue conflict. In CA1, subsets of place cells may preferentially align either with proximal cues or with distal landmarks, resulting in split representations of space^16,23^. By comparison, CA3 place cells are more strongly controlled by proximal cues^16^, highlighting differential cue weighting even across hippocampal subfields.

With their hexagonal firing patterns, grid cells in the medial entorhinal cortex (MEC) provide a local metric of the environment within the brain and dynamically switch between different reference frames^4,24–26^. By integrating self-motion and sensory information, grid cells are thought to contribute critically to error correction during path integration^27,28^. Consistent with this role, grid cells generally follow visual landmarks when visual and idiothetic cues are experimentally dissociated, but under conditions of pronounced conflict, grid representations can instead remain aligned to self-motion cues, suggesting flexible and context-dependent cue weighting^29^. A pressing but yet open question has remained how cue weighting of local versus distal cues determines grid cell mediated representations.

Importantly, cue weighting is not static, but adapts according to prior experience and perceived cue reliability. Grid cell firing patterns can become distorted or remapped depending on the stability of environmental relationships and previous experience within an environment^18,30,31^. Similarly, place cells in CA1 and HD cells in the ADN follow mobile landmarks unless they are perceived as unreliable^32,33^. More broadly, work on multisensory integration shows that neural systems often combine cues in a statistically principled, reliability-weighted fashion, adjusting perceptual weights when cue reliability changes with experience^6,34–38^. However, it has remained underexplored whether this process changes dynamically based on the expected cue reliability.

Here, we set out to systematically test how cue weighting of local versus distal cues affects spatial representations in the MEC. Furthermore we investigated over which time scales the perceived coherent or non-coherent cue relationships dictate subsequent spatial coding.

## Results

### Prior cue coherence predicts grid cell stability in darkness

To test how prior experience dynamically influences grid cell firing, we performed electrophysiological recordings in 12 mice implanted with silicon probes targeting the superficial layers of MEC. The experimental setup consisted of a rotating arena positioned in a corner of the recording room, allowing the adjacent walls and the room environment to serve as distal cues (**Figure 1 A,E**). The recording protocol entailed a 30min baseline random-foraging trial in light, followed by ten 2min dark intervals interleaved with nine 2min light intervals, and a final 30min baseline light trial. Mice were assigned to one of the two groups, henceforth termed “stable” or “rotation”. In the stable group, animals were exposed to darkness after experiencing no alterations in the surrounding environment during light trials, i.e. fixed relationship between arena and distal room cues (**Figure 1 A**). In contrast, mice in the rotation group were subjected during the 2min light intervals to a changing environment, i.e. the arena was rotated by a random angle (90–180°) thereby introducing a mismatch between arena and distal cues. (**Figure 1 E**). Each mouse completed six sessions, which are referred to as “habituation phase” in the following.

**Figure 1:**
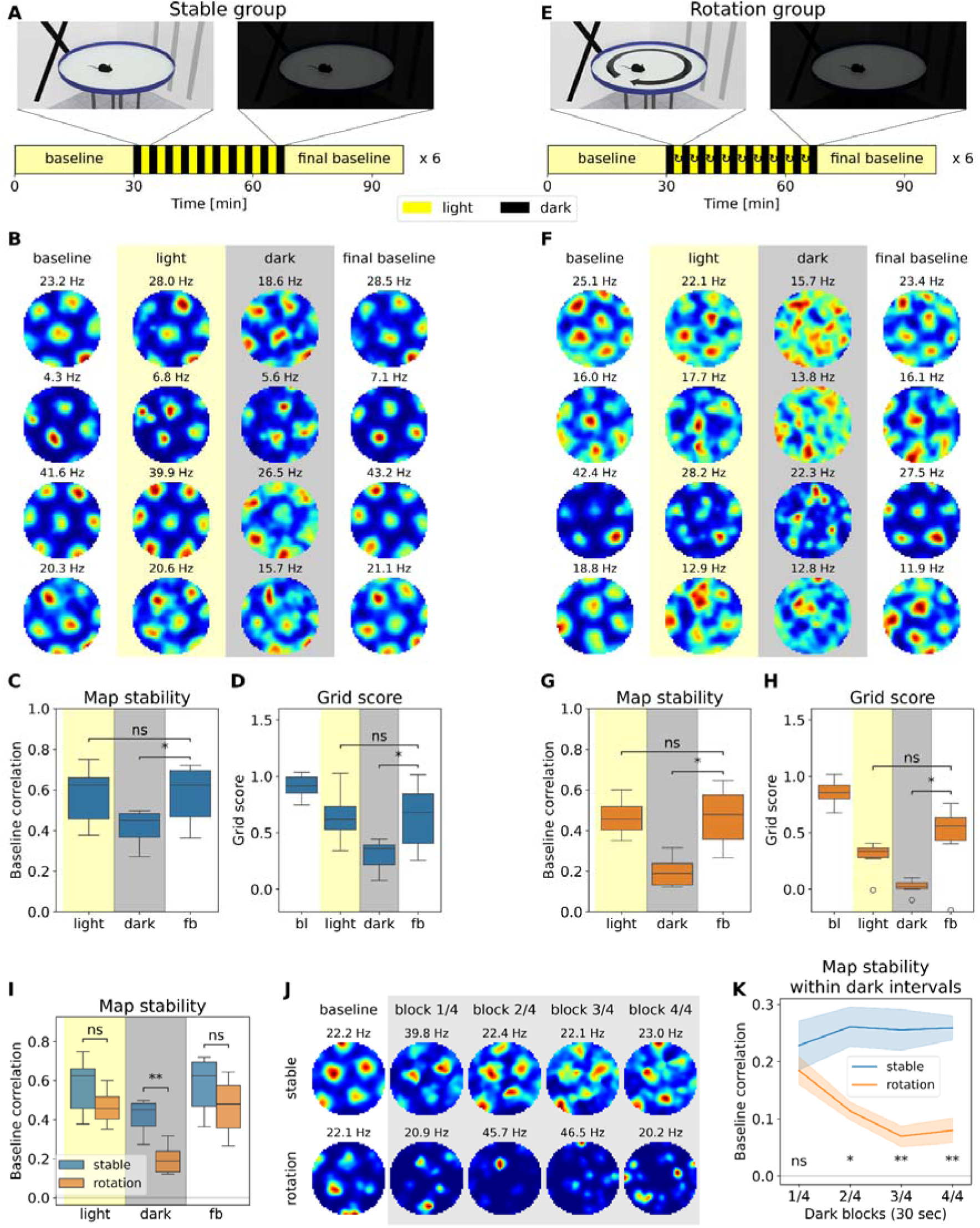
Spatial coherence between local and distal cues determines grid cell stability in subsequent darkness. **A**Top panel: Schematic of the recording setup during light and dark intervals. The black cross and the silver stripes on the walls represent the distal room cues. Bottom panel: Recording protocol to which mice of the stable group were exposed: A 30min baseline trial of random foraging under light conditions is followed by ten 2min dark intervals, interleaved with nine 2min light intervals. The protocol entails a final 30min random-foraging trial in light. Each animal was exposed to six sessions on consecutive days. **B** Example firing rate maps of four representative grid cells from the stable group, plotted for the baseline trial, the concatenated light intervals (“light”), the concatenated dark intervals (“dark”) and the final baseline. The peak firing rate is indicated above each firing rate map. The grid scores are as follows: <u>cell1</u>: 1.37, 1.04, 1.25, 1.36; <u>cell2</u>: 1.25, 1.16, 1.25, 1.08; <u>cell3</u>: 1.39, 1.38, 0.92, 1.36; <u>cell4</u>: 1.32, 1.02, 1.07, 1.28 **C** Map stability of grid cells in the stable group in light, dark and the final baseline (fb) compared to the baseline trial. Map stability is defined as the correlation of the firing rate map in the baseline trial with that in the subsequent trials. Data is grouped by mouse. Wilcoxon signed-rank test, one-sided, statistical unit: mouse: n=6; light vs fb: p=0.5781, W=11; dark vs fb: p=0.01562, W=0. **D** Grid scores of grid cells in the stable group for the baseline trial (bl), light, dark and the final baseline. The data is grouped by mouse. Wilcoxon signed-rank test, one-sided, statistical unit: mouse: n=6; light vs fb: p=0.5781, W=11; dark vs fb: p=0.01562, W=0. **E** Top panel: Schematic of the recording setup during light and dark intervals. The black cross and the silver stripes on the walls represent the distal room cues. Bottom panel: Recording protocol to which mice of the stable group were exposed: The protocol is similar to that of the stable group, but the arena is rotated by a random angle (90°-180°) in each light interval. Each animal was exposed to six sessions on consecutive days. **F** Same as in B, but for the rotation group. The grid scores are as follows: <u>cell1</u>: 1.09, 0.99, –0.02, 0.66; <u>cell2</u>: 1.22, 0.62, –0.01, 1.19; <u>cell3</u>: 1.20, 1.01, –0.11, 1.29; <u>cell4</u>: 1.42, 1.09, –0.33, 1.52 **G** Same as in C, but for the rotation group. Wilcoxon signed-rank test, one-sided, statistical unit: mouse: n=6; light vs fb: p=0.5781, W=11; dark vs fb: p=0.01562, W=0. **H** Same as in D, but for the rotation group. Wilcoxon signed-rank test, one-sided, statistical unit: mouse: n=6; light vs fb: p=0.07812, W=3; dark vs fb: p=0.03125, W=1. **I** Map stability of grid cells in light, dark and final baseline comparing the stable and the rotation group. Mann-Whitney-Wilcoxon test, two-sided, statistical unit: mouse: n(stable)=6, n(rotation)=6; light: p=0.1797, U=27; dark: p=0.004329, U=35; fb: p=0.2403, U=26. **J** Example firing rate maps of a grid cell in the baseline trial and in darkness for the stable group (top panel) and the rotation group (bottom panel). Each 2min dark interval is split into four 30sec blocks, which are then concatenated across intervals. the peak firing rate is indicated above each firing rate map. The grid scores for the baseline firing rate maps are as follows: <u>cell1</u>: 1.26, <u>cell2</u>: 1.20. The map stability values for the dark blocks are as follows: <u>cell1</u>: 0.74, 0.82, 0.64, 0.64; <u>cell2</u>: 0.25, 0.16, 0.01, – 0.02 **K** Map stability of grid cells during the 4 blocks of darkness compared between the stable and the rotation group. Mann-Whitney-Wilcoxon test, two-sided, statistical unit: mouse: n(stable)=6, n(rotation)=6; block 1/4: p=0.3095, U=25; block 2/4: p=0.01515, U=33; block 3/4: p=0.002165, U=36; block 4/4: p=0.002165, U=36.

Grid cells were identified using standard grid-score criteria and shuffled significance thresholds. The number of grid cells and their baseline grid scores were comparable between the two groups (**Supplementary Figure S1**). Grid cell firing rates and running speed also remained similar between groups and light-dark conditions (**Supplementary Figure S2**), indicating comparable behavioral engagement.

Grid cells in the stable group maintained stable firing patterns in both light and dark conditions (**Figure 1 B**), although the grid score and map stability relative to the baseline trial were slightly reduced (**Figure 1 C,D**). In contrast, in the rotation group, there was a near-complete loss of the grid structure in darkness (**Figure 1 F-H**). Accordingly, grid cell stability in darkness was significantly lower in the rotation group compared to the stable group (**Figure 1 I**).

We next examined the temporal dynamics of map stability within each 2min dark interval. To this end, each interval was divided into four 30sec blocks, which were then concatenated across intervals (**Figure 1 J**). In the stable group, the grid structure remained by and large constant across all blocks. In contrast, in the rotation group, there was a rapid decline in map stability, reaching near-chance level within approximately 60 seconds (**Figure 1 J,K**). The fitted decay time constant was in the order of 10–15 seconds.

We conclude that grid cell stability in darkness, based on grid score and map stability, critically depends on prior experience of coherent relationships between arena and distal cues.

### Grid cells preferentially anchor to distal cues

During the light intervals, when using the rotation protocol, the arena was rotated by a random angle, thereby eliminating a consistent relationship between local cues on the arena and distal room cues. To determine the reference frame of grid cell activity under cue-mismatch conditions, we computed and compared firing rate maps for both room– and arena-centered coordinates. Grid cells exhibited clear anchoring to the room reference frame in light (**Figure 2 A**). Specifically, grid scores were markedly reduced when rate maps were computed using the arena as reference frame, whereas maps referenced to the room frame preserved robust grid structure (**Figure 2 B)**. In contrast, during darkness, grid cell activity was not anchored to either the room reference frame (confirming that distal cues were not available to the mice) or the arena reference frame. These results demonstrate that grid cells preferentially encode space relative to distal cues.

**Figure 2:**
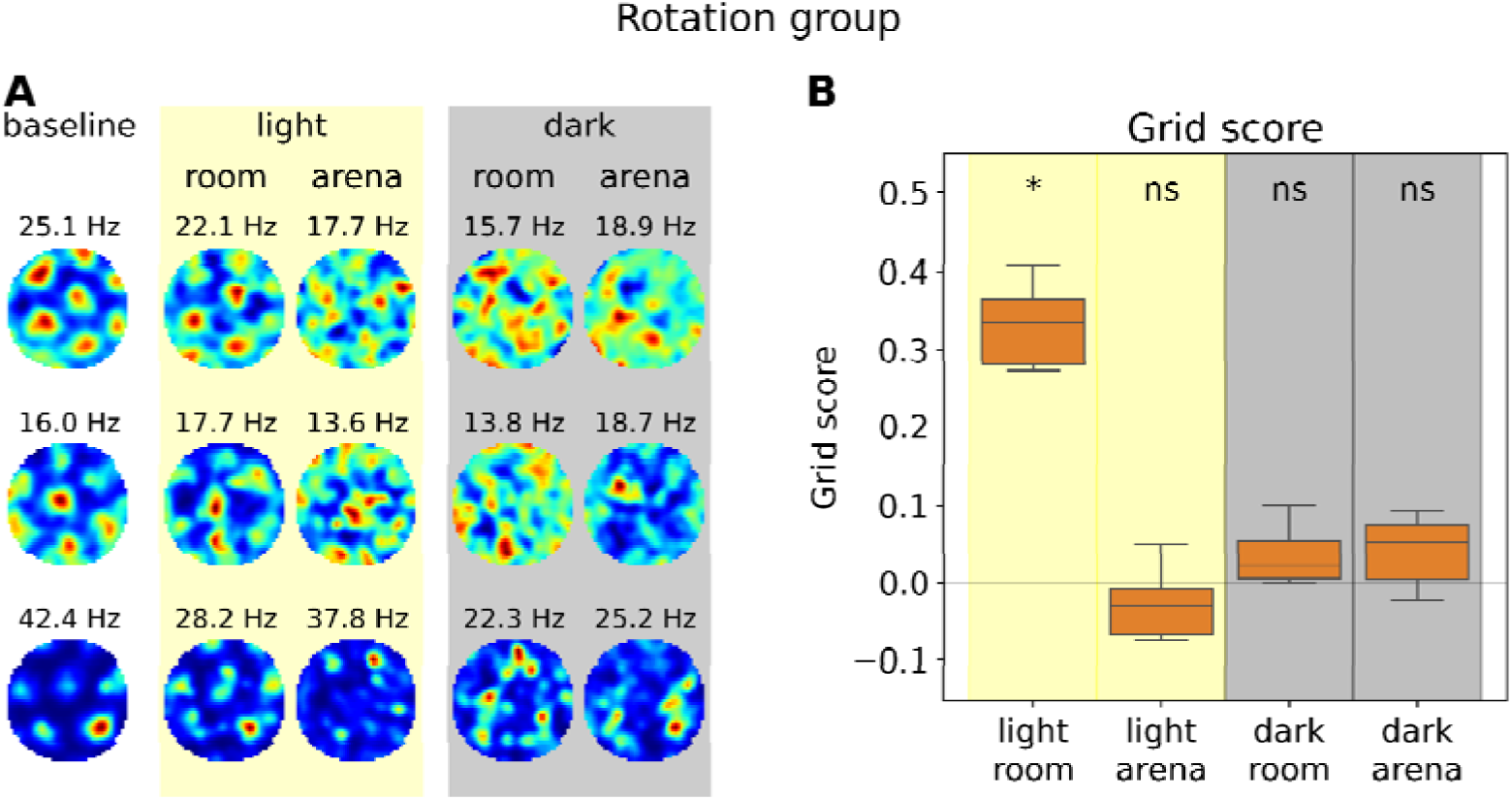
In a cue-conflict scenario, grid cells rely preferentially on distal cues. **A** Example firing rate maps of three representative grid cells from the rotation group for baseline, light and dark. For light and dark, the firing rate maps in the room reference frame as well as in the arena reference frame are shown. Note that the room and arena reference frame are not identical in light conditions due to rotation of the arena, and in darkness, as the orientation of the arena changes after each rotation in light. The peak firing rate is indicated above each firing rate map. The grid scores are as follows: <u>cell1</u>: 1.09; 0.99, –0.02, –0.02, –0.02; <u>cell2</u>: 1.22, 0.62, –0.06, –0.01, –0.17; <u>cell3</u>: 1.20, 1.01, 0.04, –0.11, –0.30 **B** Grid score of grid cells from the rotation group for light and dark in the room and arena reference frame. Wilcoxon signed-rank test against zero, one-sided, statistical unit: mouse: n=6; light room: p=0.03125, W=20; light arena: p=0.9531, W=3; dark room: p=0.2188, W=15; dark arena: p=0.07.813, W=18.

### Repeatedly experienced cue coherence in light gradually improves grid stability in darkness

To assess whether grid stability changed over the course of a session, we compared early versus late dark intervals within each session. In the stable group, map stability remained high throughout the sequence of dark intervals and did not differ significantly between the first and last five intervals, although there was a slight decrease in grid score. In the rotation group, map stability was already low in the first dark intervals and remained low in the later intervals, without significant within-session change (**Figure 3 A-B**). Across sessions, however, the stable group showed a progressive increase in grid score and a similar trend in map stability in darkness, whereas the rotation group remained at consistently low, near-chance levels (**Figure 3 C-D**). These results indicate that repeated exposure to coherent proximal and distal cues in light led to a gradual enhancement of grid cell stability in darkness across days.

**Figure 3:**
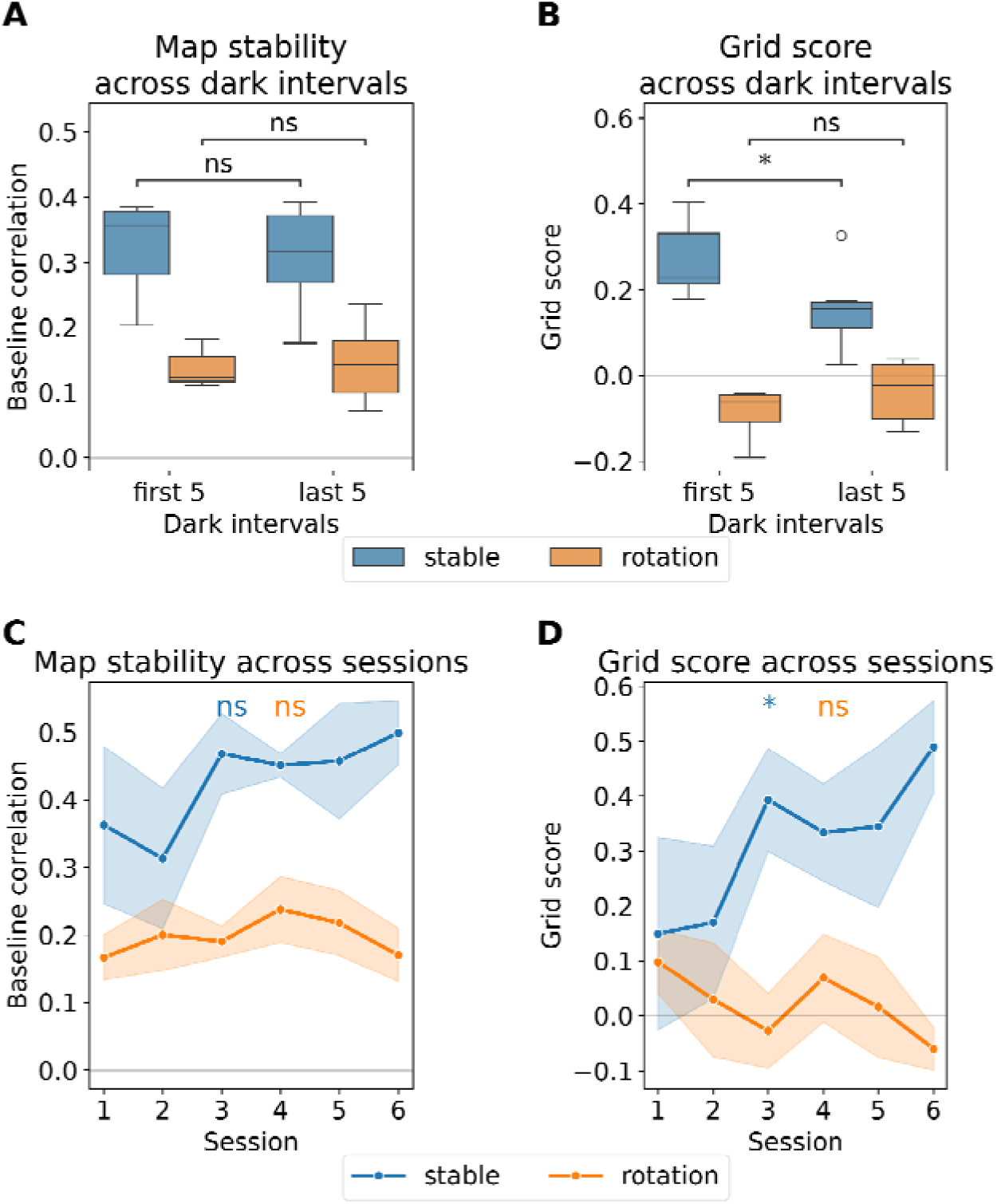
Repeated exposure to coherent proximal and distal cues in light progressively increases grid cell stability across sessions, but not within sessions. **A** Map stability of grid cells from the stable group and from the rotation group compared between the first 5 and the last 5 dark intervals. Wilcoxon signed-rank test, two-sided, statistical unit: mouse: stable: n=6, p=0.4375, W=6; rotation: n=6, p=0.6875, W=8. **B** Same as in A, but the grid score is shown. Wilcoxon signed-rank test, two-sided, statistical unit: mouse: stable: n=6, p=0.03125, W=0; rotation: n=6, p=0.2188, W=4. **C** Map stability of grid cells across the six recording sessions comparing the stable and the rotation group. The mean with the standard error are shown. Linear mixed-effects model (random intercept for subject): statistical unit: mouse; stable: n=6, β=0.032/session, p=0.088; rotation: n=6, β=0.001/session, p = 0.870. **D** Same as in C, but the grid score is shown. Linear mixed-effects model (random intercept for subject): statistical unit: mouse; stable: n=6, β=0.062/session, p=0.039; rotation: n=6, β=-0.021/session, p=0.238.

### Prior cue coherence in light also determines HD cell stability in darkness

To test whether this dependence on prior cue coherence pertains not only to grid cells, but also to other spatially-tuned cell types in the MEC, we analyzed the stability of head direction (HD) cells (**Figure 4 A,B**). The number of HD cells and their baseline HD scores were comparable between the two groups (**Supplementary Figure S3**). In the stable group, HD score and tuning stability (quantified as baseline correlation) remained high throughout dark intervals, consistent with preserved directional representations when local and distal cues had been coherently aligned in prior light intervals (**Figure 4 C,D**). In contrast, the rotation group exhibited a marked reduction in both HD tuning stability and HD score during darkness (the latter statistically not significant, but with a strong tendency), closely paralleling the disruption in grid cells after repeated experience of cue mismatch in light (**Figure 4 C,D**). This was evident also at finer temporal resolution (**Figure 4 E**). Together, these findings imply that the impact of prior cue coherence is not limited to spatial metric representations, but extends to directional coding, suggesting a shared, history-dependent mechanism of cue integration across entorhinal cell types.

**Figure 4:**
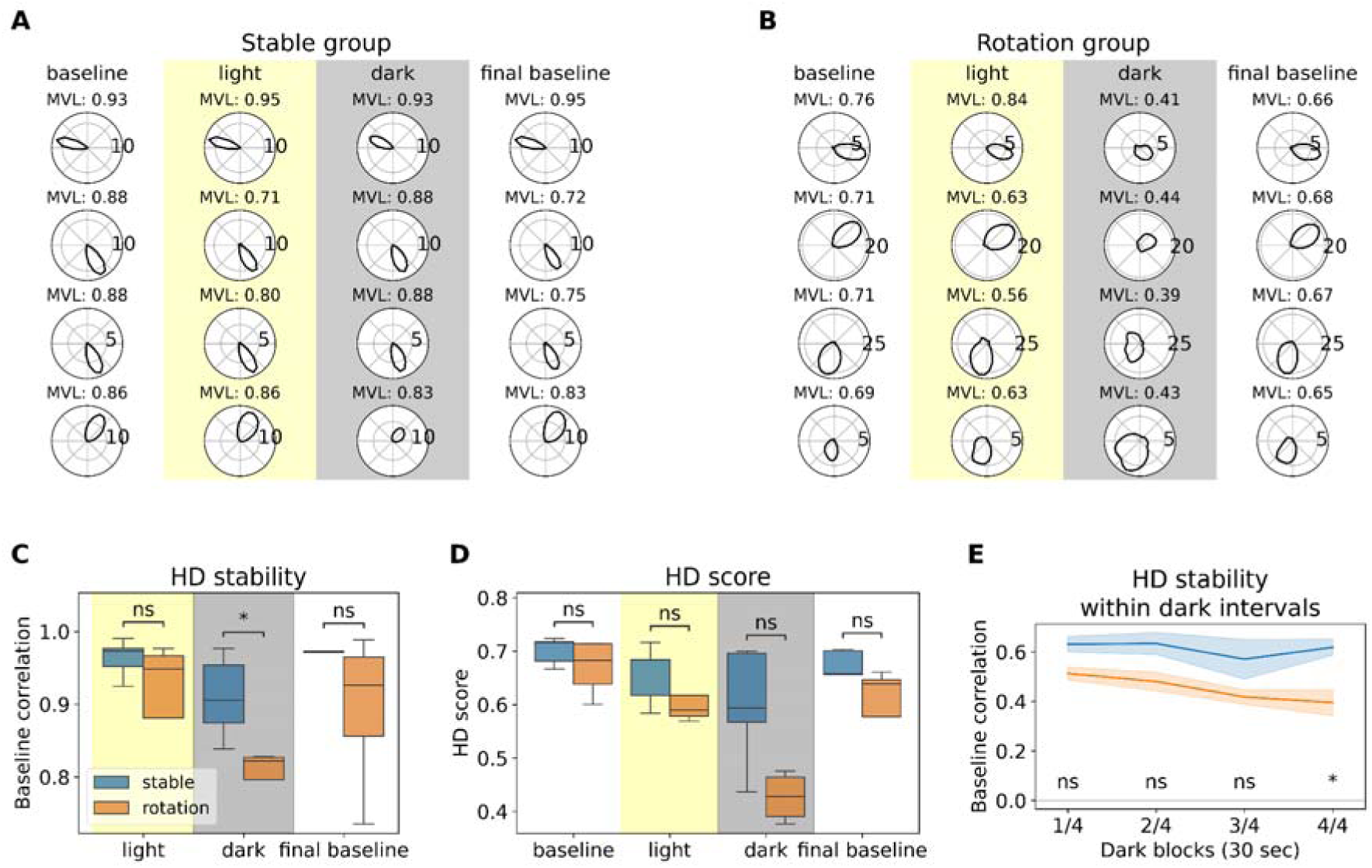
Head-direction (HD) cell stability in darkness is disrupted in mice previously exposed to cue mismatch in light. **A** Example polar plots of four representative HD cells from the stable group for baseline, light, dark and final baseline. The HD score (mean vector length, MVL) is indicated above each polar plot. **B** Same as in A, but for the rotation group. **C** HD stability in light, dark and in the final baseline compared between the stable and the rotation group. HD stability in a given condition is defined as the correlation of the HD tuning curve for the baseline trial and that of this condition. Mann-Whitney-Wilcoxon test, two-sided, statistical unit: mouse: n(stable)=5, n(rotation)=4; light: p=0.5556, U=13; dark: p=0.01587, U=20; fb: p=0.4127, U=14. **D** Same as in C, but the HD score is shown. The legend from panel C applies. Mann-Whitney-Wilcoxon test, two-sided, statistical unit: mouse: n(stable)=5, n(rotation)=4; baseline: p=0.1905, U=16; light: p=0.1111, U=17; dark: p=0.06349, U=18; fb: p=0.2857, U=15. **E** HD stability within darkness compared between the stable and rotation group. Each 2min dark interval is split into four 30sec blocks, which are then concatenated across intervals. The legend from panel C applies. Mann-Whitney-Wilcoxon test, two-sided, statistical unit: mouse: n(stable)=5, n(rotation)=4; block 1/4: p=0.06349, U=18; block 2/4: p=0.06349, U=18; block 3/4: p=0.1905, U=16; block 4/4: p=0.01587, U=20.

### Cue coherence in light is a prerequisite for the switch from distal to proximal cues in subsequent darkness

In the rotation group, grid stability declined rapidly, whereas in the stable group a robust grid structure was maintained throughout darkness (**Figure 1 K**). This raises the question of how grid stability is sustained for so long in the absence of visual input. To address this question, we asked whether grid cells could anchor to the arena using local cues on the arena. Thus, after completion of the 6 sessions, the mice underwent a test session, during which the arena was slowly and continuously rotated in darkness (at a speed of 1.6°/sec) below the vestibular detection threshold^39^. The protocol was identical for both groups (i.e. no rotation in light), so that any difference in grid cell coding would reflect prior learning history (**Figure 5 A,C**). Of note, there was a significant difference between the stable and the rotation group when considering map stability and grid score in darkness in the arena reference frame (**Figure 5 B,D-F**). The difference in map stability persisted throughout the session (**Figure 5 G**).

**Figure 5:**
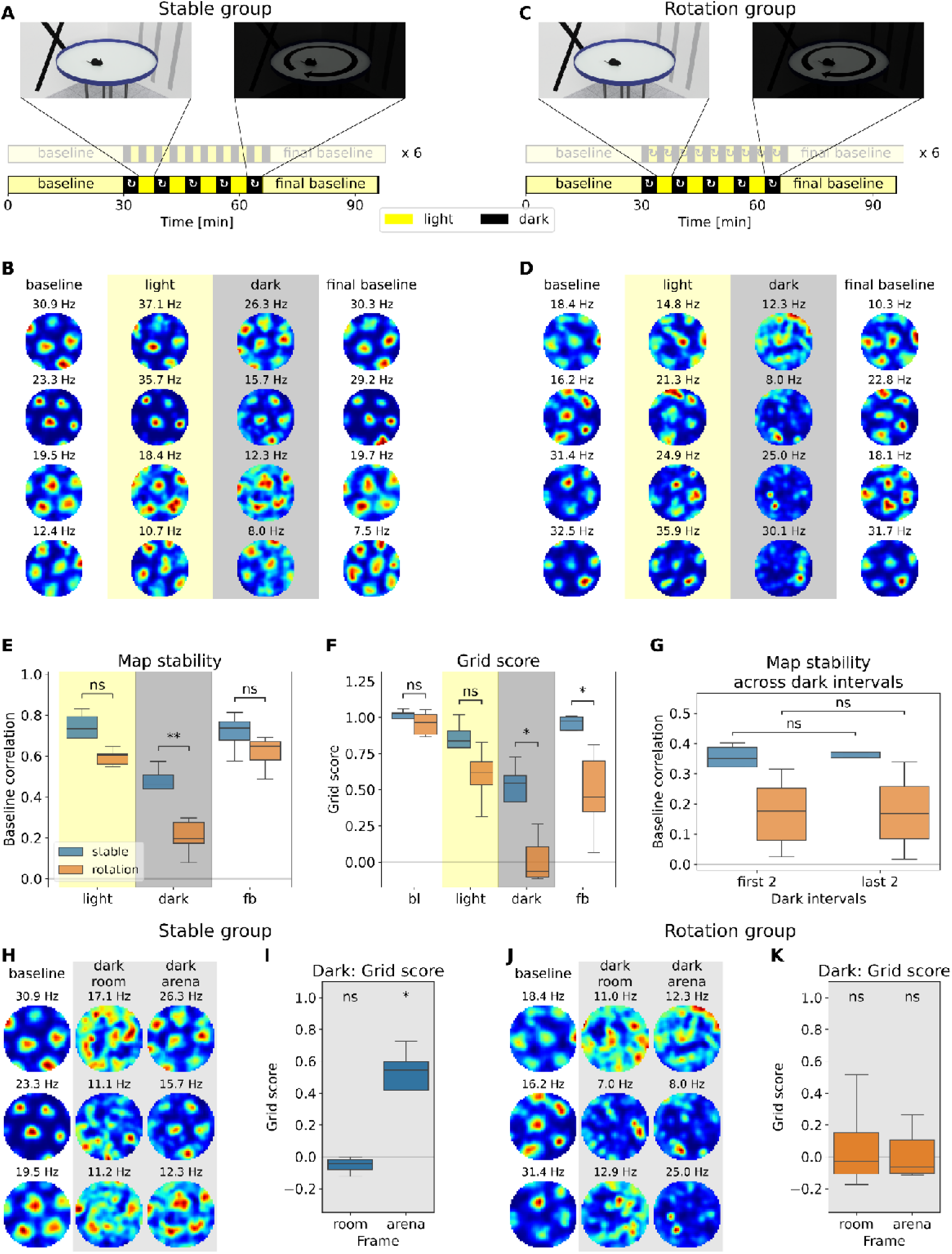
Grid cells anchor to the rotating arena in darkness when previous light experience of proximal and distal cues was coherent. **A** Recording protocol for the stable group: After the habituation phase (six sessions without rotation in light, greyed-out protocol), the mice are exposed to the following slow rotation protocol: A 30min baseline random-foraging trial is followed by five 4min dark intervals, interleaved with three 4min light intervals. During each of the dark intervals, the arena is rotated by 360°. The protocol concludes with a final 30min random-foraging trial in light. **B** Example firing rate maps of four representative grid cells from the stable group, plotted for the baseline trial, the concatenated light intervals, the concatenated dark intervals and the final baseline trial in the arena reference frame. The peak firing rate is indicated above each firing rate map. The grid scores are as follows: <u>cell1</u>: 1.38, 1.19, 1.27, 1.20; <u>cell2</u>: 1.21, 1.28, 1.14, 1.25; <u>cell3</u>: 1.41, 1.33, 1.12, 1.18; <u>cell4</u>: 1.25, 0.91, 1.16, 1.16 **C** Recording protocol for the rotation group: After the habituation phase (6 sessions with rotation in light, greyed out), the mice are exposed to the same slow rotation protocol as the stable group. **D** Same as in B, but for the rotation group. The grid scores are as follows: <u>cell1</u>: 1.22, 1.08, –0.08, 1.37; <u>cell2</u>: 1.13, 0.54, –0.30, 0.80; <u>cell3</u>: 1.27, 1.20, 0.33, 1.10; <u>cell4</u>: 1.23, 0.96, –0.15, 1.20 **E** Map stability of grid cells in light, dark and final baseline (fb) compared between the stable and the rotation group. Mann-Whitney-Wilcoxon test, two-sided, statistical unit: mouse: n(stable=5, n(rotation)=6; light: p=0.1255, U=24; dark: p=0.008658, U=29; fb: p=0.08225, U=25. **F** Grid scores of grid cells in baseline (bl), light, dark and final baseline compared between the stable and the rotation group. The legend from panel E applies. Mann-Whitney-Wilcoxon test, two-sided, statistical unit: mouse: bl: p=0.4286, U=20; light: p=0.1255, U=24; dark: p=0.01732, U=28; fb: p=0.03030, U=27. **G** Map stability of grid cells from the stable group and from the rotation group compared between the first two and the last two dark intervals. The legend from panel E applies. Wilcoxon signed-rank test, two-sided, statistical unit: mouse: stable: n=5, p=0.6250, W=5; rotation: n=6, p=1.000, W=10. **H** Example firing rate maps of three representative grid cells in the baseline trial and in the room and arena reference frame in darkness for the stable group. The peak firing rate is indicated above each firing rate map. The grid scores are as follows: <u>cell1</u>: 1.38, –0.34, 1.27; <u>cell2</u>: 1.21, –0.19, 1.14; <u>cell3</u>: 1.41, 0.13, 1.12 **I** Grid score of grid cells from the stable group in the room and arena reference frame in darkness. The data is grouped by mouse. Wilcoxon signed-rank test, one-sided, statistical unit: mouse: n=5; room: p=1.000, W=0; arena: p=0.03125, W=15. **J** Same as in H, but for the rotation group. The grid scores are as follows: <u>cell1</u>: 1.22, 0.28, –0.08; <u>cell2</u>: 1.13, –0.02, –0.30; <u>cell3</u>: 1.27, –0.31, 0.33 **K** Same as in I, but for the rotation group. Wilcoxon signed-rank test, one-sided, statistical unit: mouse: n=6; room: p=0.5000, W=11; arena: p=0.5000, W=11.

In the stable group, grid cells retained a clear grid pattern in darkness when firing rate maps were computed in the arena reference frame (**Figure 5 B,H**), whereas grid scores in the room frame were at chance levels, indicating that the internal map tracked the arena rather than the room (**Figure 5 H,I**). In contrast, grid cells in the rotation group did not show reliable anchoring to either frame (**Figure 5 D,J,K)**, indicating that repeated cue mismatch precluded the use of the arena as a stable proximal reference. These findings demonstrate that prior experience of coherent cue relationships is necessary to deploy proximal cue information to support grid stability even when visual input is absent.

### Brief exposure to cue incoherence precludes the switch from distal to local cues in darkness

Given that grid cell activity differed markedly as a function of prior experience, we wondered whether the differential effects in the two groups could be reversed by exposing the mice to the protocol of the opposite group. Thus, mice from the stable group underwent the rotation protocol and vice versa (**Figure 6 A,E**). When animals from the stable group were switched to the rotation protocol, grid stability in darkness dropped sharply, indicating that previously learned coherent cue associations were rapidly disrupted once cue set relationships became incoherent (**Figure 6 B-D**). In contrast, grid cells in mice of the rotation group, when switched to the stable protocol, exhibited substantial inter-individual variability in both grid score and map stability, suggesting that grid cells only in a subset of animals were anchored to local cues despite the restored coherence between proximal and distal cues (**Figure 6 F-H**). Accordingly, grid stability in the stable-to-rotation condition dropped to levels comparable to those observed in the rotation group exposed to the rotation protocol from the beginning, whereas the reverse transition did not occur at a comparable time scale (**Figure 6 I**).

**Figure 6:**
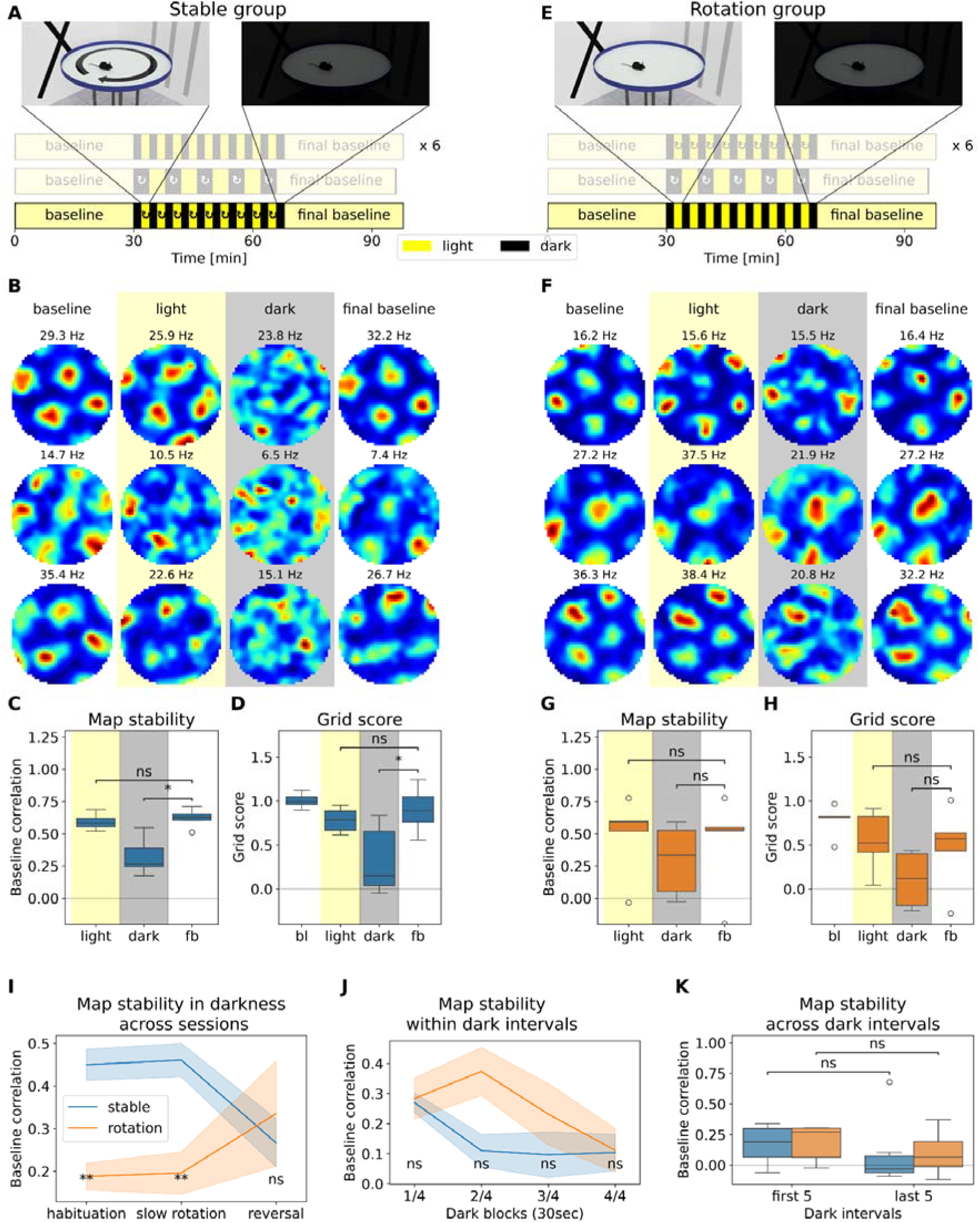
Stabilization and destabilization of grid cell patterns occur at different time scales. **A** Recording protocol for the stable group: After the habituation phase (six sessions without rotation in light, greyed-out) and the slow rotation in darkness (greyed-out), the mice are exposed to the reversed-habituation protocol (bottom panel): A 30min baseline random-foraging trial is followed by ten 2min dark intervals, interleaved with nine 2min light intervals. The arena is rotated by a random angle (90°-180°) in each dark interval. The protocol concludes with a final 30min random-foraging trial in light. This protocol is thus identical to the habituation protocol of the rotation group. **B** Example firing rate maps of three representative grid cells from the stable group, plotted for the baseline trial, the concatenated light intervals, the concatenated dark intervals and the final baseline of the reversed-habituation session. The peak firing rate is indicated above each firing rate map. The grid scores are as follows: <u>cell1</u>: 1.19, 0.97, –0.18, 1.33; <u>cell2</u>: 1.04, 0.72, –0.22, 0.87; <u>cell3</u>: 0.90, 0.92, 0.84, 0.55 **C** Map stability of grid cells in the stable group in light, dark and the final baseline (fb) compared with the baseline trial. Data is grouped by mouse. Wilcoxon signed-rank test, one-sided, statistical unit: mouse: n=6; light vs fb: p=0.2188, W=6; dark vs fb: p=0.01562, W=0. **D** Grid scores of grid cells in the stable group for the baseline trial (bl), light, dark and the final baseline. The data is grouped by mouse. Wilcoxon signed-rank test, one-sided, statistical unit: mouse: n=6; light vs fb: p=0.1562, W=5; dark vs fb: p=0.03125, W=1. **E** Recording protocol for the rotation group: After the habituation phase (six sessions with rotation in light, greyed-out) and the slow rotation in darkness (greyed-out), the mice are exposed to the reversed-habituation protocol (bottom panel): A 30min baseline random-foraging trial is followed by ten 2min dark intervals, interleaved with nine 2min light intervals. The protocol concludes with a final 30min random-foraging trial in light. This protocol is thus identical to the habituation protocol of the stable group. **F** Same as in B, but for the rotation group. The grid scores are as follows: <u>cell1</u>: 1.30, 1.32, 0.55, 1.38; <u>cell2</u>: 1.00, 0.96, 1.20, 1.06; <u>cell3</u>: 1.06, 0.97, 0.31, 0.95 **G** Same as in C, but for the rotation group. Wilcoxon signed-rank test, one-sided, statistical unit: mouse: n=5; light vs fb: p=0.9062, W=12; dark vs fb: p=0.1562, W=3. **H** Same as in D, but for the rotation group. Wilcoxon signed-rank test, one-sided, statistical unit: mouse: n=5; light vs fb: p=0.7812, W=10; dark vs fb: p=0.1562, W=3. **I** Map stability of grid cells in dark in the habituation phase, the slow rotation protocol and the reversed-habituation (“reversal”) for the stable and the rotation group. For the habituation phase, the map stability is pooled across all six sessions. Mann-Whitney-Wilcoxon test, two-sided, statistical unit: mouse: n(stable)=5, n(rotation)=5; habituation: p=0.004329, U=35; slow rotation: p=0.008658, U=29; reversal: p=1.000, U=15. **J** Map stability of grid cells during the four blocks of darkness in the reversed-habituation protocol compared between the stable and the rotation group. The legend from panel I applies. Mann-Whitney-Wilcoxon test, two-sided, statistical unit: mouse: n(stable)=6, n(rotation)=5; block 1/4: p=0.9307, U=16; block 2/4: p=0.3290, U=9; block 3/4: p=0.6623, U=12; block 4/4: p=0.9307, U=14. **K** Map stability of grid cells from the stable group and from the rotation group compared between the first 5 and the last 5 dark intervals of the reversed-rotation protocol. The legend from panel I applies. Wilcoxon signed-rank test, two-sided, statistical unit: mouse: stable: n=6, p=0.6875, W=8; rotation: n=5, p=0.4375, W=4.

At finer temporal resolution, grid stability in the stable-to-rotation condition declined within the first 30sec of darkness, closely matching the rapid decay observed during initial exposure to cue mismatch (**Figure 6 J**, **Figure 1 K**). Conversely, in the rotation-to-stable condition, grid cells maintained relatively high stability only during the first half of each dark interval, despite restored cue coherence (**Figure 6 J**). Notably, no significant within-session changes were observed when comparing early and late dark intervals in either group (**Figure 6 K**). In sum, these findings indicate that a single experience of incoherent local and distal cues can abolish the use of proximal cues for spatial coding in darkness, whereas recovery following reintroduction of coherent cues is slower and incomplete.

## Discussion

Spatial navigation relies on the integration of multiple sources of information, including sensory landmarks and self-motion cues, to estimate position and guide movement. The contribution of each cue depends on its salience and perceived reliability, both of which are shaped through experience^7–14^. Our findings reveal an additional level of flexibility in this process: Although the relationship between cue sets is learned over days, a single experience of cue mismatch sufficed to alter the inferred reliability of the cues. Specifically, when mice experienced a stable relationship between distal room cues and local arena cues in light across days, the grid pattern of grid cells stabilized in darkness, while a single exposure to cue mismatch abolished the learned dependence on local cues. Strikingly, the reverse regime switch (i.e., the experience of a coherent cue set relationship after repeated exposure to cue mismatch) led only to an incomplete recovery with great inter-mouse variability. We conclude that inferred cue reliability depends not only on how often and how recently a cue relationship is experienced, but also on whether the expected relationship is confirmed or violated.

The behavioral paradigm allows the dissection of four scenarios in which distinct spatial representations reflect differential strategies of cue use. First, during light sessions involving a mismatch between distal room cues and local arena cues, grid cells consistently remained anchored to the room reference frame, demonstrating a clear preference for distal visual landmarks over local cues. Second, grid representations in darkness degraded within tens of seconds when there was a cue mismatch in prior light experience. This temporal profile is consistent with a navigation strategy that relies predominantly on path integration without error correction by local sensory information^40^. Third, grid firing remained stable throughout darkness in mice that had experienced only stable cue relationships during preceding light sessions. The absence of progressive degradation argues against path integration alone sustaining the representation over these timescales. Instead, slowly rotating the arena during darkness demonstrated that grid cells remained anchored to the arena rather than to the room reference frame. Fourth, the findings related to scenario three further imply that during preceding light intervals with coherent cue relationships, distal and local cues were integrated into a common spatial representation. Because both reference frames were aligned, the integration supported robust and stable grid firing.

The issue of whether path integration plays a role in this paradigm merits further consideration. We propose that in the rotation condition in light, when there is a mismatch between distal and local cues, and grid cells rely on a distal reference frame, not only local cues become devaluated, but also self-motion cues. Thus, we can not exclude the possibility that the devaluation of self-motion cues also contributes to the disruption of grid cell firing in darkness. This in turn raises the question to which extent local cues vs. self-motion cues are used to maintain a stable grid pattern in darkness following stable light conditions. We favor a dominant role of local cues, since there was no sign of error accumulation over recording times of up to four minutes, indicating that path integration relying on self-motion only is unlikely to sustain a stable grid map under these conditions.

The cooperative integration of distal and local information argues against a simple associative overshadowing mechanism, in which the dominant cue set prevents learning about weaker concurrent cues^11,15^. Despite preferential anchoring to distal landmarks in the presence of cue mismatch, local arena cues were nevertheless encoded and subsequently supported stable spatial representations in darkness. Rather than suppressing learning, coherent cue relationships therefore appear to promote the establishment of multiple complementary spatial anchors that can be flexibly recruited when environmental conditions change.

Importantly, within such a framework, one can easily reconcile apparently conflicting results, including work from this laboratory, regarding grid cell stability in darkness. Thus, studies in which the relationship between room and arena cues remained stable during interleaved light sessions consistently reported preserved grid firing in darkness, albeit with modestly reduced grid scores compared with light^24,41,42^, potentially reflecting the greater precision afforded by visual landmarks relative to haptic or olfactory arena cues. In contrast, studies in which the dark trial was recorded first or distal cues were experimentally manipulated before darkness reported degraded grid representations^40,43,44^. Together, these findings suggest that maintaining a coherent relationship between distal and local cues before visual input is removed is a prerequisite for preserving stable grid representations in darkness.

Head direction cells exhibited a similar dependence on prior cue coherence. Following exposure to cue conflict, MEC HD cells displayed reduced directional stability in darkness, whereas directional tuning remained robust after stable cue experience. These observations reinforce the close functional coupling between the HD and grid cell systems^45^ and likewise provide a parsimonious explanation for previously inconsistent findings across studies^40,42^ (but see Chen et al.^43^).

Unlike grid and HD cells, hippocampal place cells preferentially follow local cues, particularly in CA3, whereas CA1 populations can simultaneously represent both local and distal reference frames^16,23^. It is therefore plausible that hippocampal representations contribute to the ability of grid cells to anchor to local cues when distal visual landmarks are unavailable.

The immediate neglect of local cues after a single exposure to cue mismatch indicates a low threshold for detecting unreliability. This is reminiscent of how hippocampal place cells and thalamic HD cells rapidly downweight landmarks that violate the prevailing spatial reference frame^32,38,46^. In contrast, a single experience of cue coherence is not sufficient for anchoring to a cue set that was previously deemed unreliable. Such an inference is in accordance with studies that demonstrate that place cells and HD cells retain a bias toward a previously established reference frame^33,46,47^. Together, these findings point toward a bias for conservative cue-usage strategies, favoring underuse of potentially helpful cues over reliance on misleading ones.

More broadly, our findings suggest that cue weighting within the spatial navigation system is not determined solely by long-term learning but can be rapidly reconfigured by a single experience of cue mismatch. Such asymmetric updating would allow rapid rejection of misleading cues while requiring stronger evidence before previously unreliable cues regain influence, thereby promoting robust spatial representations in changing environments. This mechanism could represent a general computational principle by which the brain continuously balances stability against flexibility during spatial navigation. In sum, here we demonstrate that experience dynamically alters the reliability assigned to different spatial cues and thereby governs the reference frame used to compute location. Of note, we recently demonstrated that grid cells operate in different reference frames during path integration^26^. Together with the findings reported here, the interesting question arises how different reference frames are selected at different time scales depending on previous experience.

## Methods

### Subjects

Adult mice (C57BL/6 wild-type, male, n=12) were housed under a 12-hour light/dark cycle with unrestricted access to food and water, except on behavioral training and recording days, when food was mildly restricted, maintaining the weight of the mouse at 85% of its initial weight. All procedures were approved by institutional and national animal care committees and complied with applicable animal welfare regulations (Regierungspräsidium Karlsruhe in compliance with the European guidelines for the care and use of laboratory animals, 86/609/EEC, license: G-236/20).

### Surgical procedures and implantations

Mice were implanted with movable silicon probes aimed at the superficial layers of medial entorhinal cortex (MEC). Surgeries were performed under isoflurane anesthesia (induction 3–4%, maintenance 0.5–1.2% in air). Animals were positioned in a stereotaxic frame, and body temperature was maintained at 37°C using a feedback-controlled heating pad. The scalp was incised, a craniotomy was made above the right MEC, and the transverse sinus was exposed. The dura was carefully removed. Silicon probes (Cambridge NeuroTech ASSY-236 H10, 64 channels) mounted on custom microdrives were positioned above the MEC, with the probe tip at 3.1 mm from the midline and 0.2 mm anterior of the transverse sinus. They were lowered 0.6–0.8 mm into the brain at an angle of 7° in the sagittal plane, with the probe tip pointing slightly in the posterior direction. The microdrive assembly was secured to the skull using bone screws and dental cement, and a ground/reference wire was attached to a screw above the cerebellum.

Post-surgery, animals received Carprofen (dose and route per institutional guidelines) for 72h and were monitored daily for signs of discomfort or infection. After at least 5–7 days of recovery, electrodes were gradually advanced into the MEC over several days while monitoring neural activity.

### Screening and targeting of MEC

Following recovery from surgery, animals underwent a screening procedure to confirm accurate targeting of the medial entorhinal cortex (MEC) and to maximize the yield of grid cells. To this end, mice were first trained to perform a random foraging task in a separate open-field environment (70 cm square enclosure) located in a different room from that where the rotating-arena setup was positioned. During the training, mice randomly foraged for food (AIN-76A Rodent Tablet 5 mg; TestDiet), which was automatically dispensed at a rate of 1-3 pellets/min, from the pellet dispenser.

For each recording, putative single units were spike-sorted, and their spatial tuning properties were assessed in the training environment. Firing-rate maps were computed, and neurons exhibiting periodic, hexagonal spatial firing patterns consistent with grid cells were identified. Based on the presence and quality of grid-like activity, as well as depth estimates from the microdrive, an anatomical estimate of the probe location relative to the superficial MEC layers was obtained.

If the target region was inferred to be located more ventrally, the silicon probe was advanced incrementally along the dorso–ventral axis between sessions. This screening-and-adjustment procedure was repeated until a stable configuration with a high yield of grid cells was achieved. Once robust grid-cell activity was observed, animals proceeded to the rotating-arena experiments described below.

### Behavioral apparatus

Recordings were performed in a circular arena (diameter: 80cm) with a homogeneous floor surface. The arena was surrounded by a low circular wall (height: 1.6cm), which acted as a proximal boundary without obscuring distal visual cues in the surrounding room. The arena was positioned in a corner of the recording room. Thus, room cues included the two adjacent walls with metal frames (south and west side), a desk with computers (north side) and a spare arena leaning against the wall (east side).

The arena was mounted on a tapered roller bearing, enabling smooth rotation by a motorized platform driven by a stepper motor. The platform’s angular position was recorded at 10Hz. Arena orientation at each time point was interpolated from these recordings, permitting transformation of the animal’s position into the arena reference frame by applying the inverse of the logged rotation angle. Distal visual cues on the room walls provided a stable global reference frame, while the arena walls and surface formed a proximal reference frame that could rotate independently. Illumination was controlled by an overhead light source operated through an Arduino-driven relay system, enabling automated transitions between full light and complete darkness.

### Tracking and data acquisition

Animal position and head direction were tracked at 50 Hz using two infrared LEDs fixed to the headstage and an overhead infrared-sensitive camera. The LED centroid provided position, and the vector between LEDs defined heading direction. The software positrack2 (https://github.com/kevin-allen/positrack2) was used.

All tracking coordinates were expressed in a common global reference frame in the setup shared across all sessions. For each time point, we stored both the global pose (`time`, `x`, `y`, `hd`) and the pose expressed in the arena frame (`x_arena`, `y_arena`, `hd_arena`) obtained by subtracting the logged arena angle from the global coordinates.

Neural signals were acquired via a headstage preamplifier connected to a multichannel recording system (RHD2000-Series Amplifier Evaluation System, Intan Technologies, sampling rate 20kHz). Spiking activity and local field potentials were digitized and recorded continuously. Synchronization between neural and behavioral data was achieved through TTL pulses: Each camera frame triggered a TTL signal recorded alongside neural data, allowing precise temporal alignment via interpolation between frame timestamps. The software ktan (https://github.com/kevin-allen/ktan) was used.

### Hardware control

Control of the entire behavioral apparatus was handled by custom Python software within a Robot Operating System (ROS) framework (https://www.ros.org). ROS nodes communicated with three key components:

- An Arduino microcontroller controlling arena rotation via stepper motor drivers.
- A dedicated Arduino–relay setup controlling the light source.
- The tracking system for frame-based TTL synchronization and logging.

ROS nodes coordinated task structure, timing, rotation commands, and event logging (e.g., arena angles, motor on/off times, light transitions). All events shared a unified time base across the ROS network and recording system, ensuring accurate synchronization for subsequent data analyses.

### Cable management system

To ensure unobstructed movement during extended recording sessions, we implemented an automated cable management system integrated with the ROS framework. A motorized actuator maintained the recording cable directly above the animal using closed-loop real-time tracking of its position. Additionally, a motorized commutator prevented cable twisting via closed-loop tracking of head direction. Both systems operated on median-filtered estimates of the last five animal poses (tracking at 50Hz) to remove tracking noise and behavioral transients.

### Recording protocols

Each recording session began with a 30 min baseline period in constant light with the arena stationary (“initial baseline”). This was followed by alternating light and dark intervals, with ten dark intervals interleaved with nine light intervals (2min each), totaling 20 min of darkness per session. The session concluded with a 30min period in constant light with the arena stationary (“final baseline”). During all parts of the recording protocol, animals foraged for scattered food rewards (AIN-76A Rodent tablets, 5 mg; TestDiet). The twelve mice were pseudo-randomly assigned to two groups (“stable” and “rotation”), with six animals per group. For the stable group, the arena remained stable throughout the session, while for the rotation group, the arena was rotated by a random angle (90°–180°, clockwise) in each 2min light interval. The rotation set in 10sec after lights on, lasted for 100sec and was followed by 10sec of stationary light before the next dark interval began. Both groups performed their respective protocol on 6 consecutive days (“habituation phase”).

On the day after the habituation phase was completed, animals of both groups completed a session following the same protocol, referred as the “slow rotation” protocol: As in the protocols described above, the session consisted of a 30min baseline trial in light conditions, followed by alternating light and dark trials and completed with a 30min baseline trial in light. However, the protocol differed from the habituation phase protocols in that five 4min dark intervals were interleaved with four 4min light intervals. During each dark interval, the arena was rotated at an angular speed below the vestibular threshold. Ten seconds after light offset, the arena began rotating, completed a full 360° revolution over the dark period and stopped 10sec before the next light onset.

On day 8, both groups were subjected to a “reversed-habituation” protocol: Animals originally assigned to the stable group were now exposed to the habituation protocol of the rotation group and vice versa.

### Electrophysiological data preprocessing

Recorded signals were band-pass filtered and spike-sorted offline using Kilosort2 (https://github.com/MouseLand/Kilosort), an automated template-matching algorithm optimized for electrophysiological recordings. Raw LFP data were filtered (300– 6,000 Hz), and clusters were manually curated in Phy (https://github.com/cortex-lab/phy) to correct sorting errors and remove noise. Unit quality was evaluated using the SpikeInterface framework (https://github.com/SpikeInterface/spikeinterface), applying the following criteria:

- Refractory period contamination ratio < 0.15
- Presence ratio ≥ 0.90
- Mean firing rate ≥ 0.5 Hz for the full session and the baseline condition Units meeting all criteria were retained for analysis.

### Spatial analysis and cell-type classification

All position and spike analyses were conducted in Python using Pynapple (https://github.com/pynapple-org/pynapple). Position data were interpolated to a common sampling grid and transformed to both room and arena reference frames.

For each interval, firing-rate maps were created by dividing spike counts by occupancy in 2cm × 2cm bins. Maps were smoothed with a Gaussian kernel (σ = 2cm). Spatial stability was quantified by correlating the firing rate map in a specfic condition with the baseline map in the same reference frame. Pearson correlation coefficients between the flattened maps provided per-cell stability scores. In analogy, HD stability in a specific condition was computed as the Pearson correlation coefficient between the HD tuning curve in this condition and the baseline trial. Grid scores were derived from the spatial autocorrelogram, defined as the difference between mean correlations at 60° and 120° vs 30°, 90°, and 150°. To calculate the HD score, a firing rate histogram per head direction was obtained using a bin size of 10° and a Gaussian kernel with a sigma of 10° for smoothing. The HD score was defined as the mean vector length of this histogram. Significance for each score was determined via 500 time-shifted spike-train surrogates. A cell was classified as a grid cell if (1) its grid score exceeded 0.45 and (2) exceeded the 95th percentile of its shuffled distribution. A cell was defined as a HD cell if (1) its HD score exceeded 0.6, (2) its HD score exceeded the 95th percentile of its shuffled distribution.

### Calculation of scores and statistical aggregation

For statistical analysis, per-session scores were obtained by averaging across all grid cells recorded from each animal in that session, yielding one grid score and one baseline correlation per animal, session, and condition (initial baseline, all light intervals, all dark intervals, final baseline). For analyses of the habituation sessions (sessions 1-6), session-level scores were further averaged per animal to obtain a single summary value per animal. The same procedure was applied to obtain per-animal HD scores and HD stability scores.

### Block binning analysis

To examine the temporal dynamics of grid cell and HD cell stability within each 2-min (120sec) dark interval, data were divided into four temporal blocks per interval: block 1 (0– 30sec), block 2 (30–60sec), block 3 (60–90sec), and block 4 (90–120sec). For analysis of grid cell firing in darkness within a recording session, all dark intervals (20min total) were concatenated and then divided into four 5min blocks per session.

### Statistical analysis

All statistical analyses were conducted in Python (SciPy). Non-parametric tests were used to avoid distributional assumptions. For within-subject comparisons (same-cell comparisons across conditions), the Wilcoxon signed-rank test was employed. For independent-group comparisons (comparisons between the stable and the rotation group), the Mann-Whitney-Wilcoxon test was used. Two-sided tests were used unless a directional hypothesis justified a one-sided test. For assessing the effect of habituation on the different groups, a linear mixed model with mice as random intercept was employed.

### Supplementary figures

**Supplementary Figure S1:**
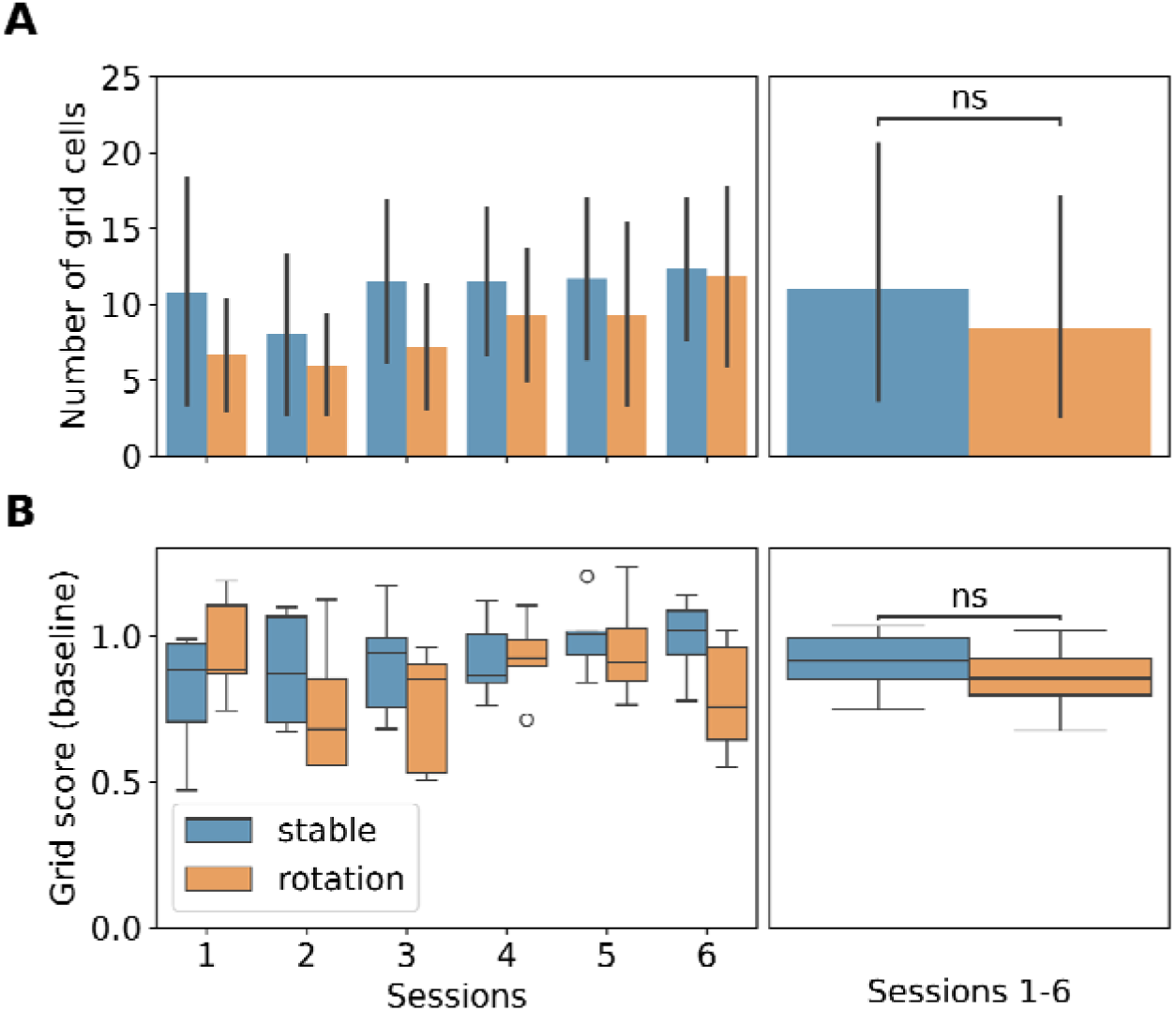
Grid cell numbers and grid scores are similar across groups. **A** Left panel: Number of grid cells in each of the 6 habituation sessions for the stable and the rotation group. Right panel: Mean number of grid cells across the 6 habituation sessions. The legend from panel B applies. Mann-Whitney-Wilcoxon test, two-sided, statistical unit: mouse: n(stable)=6, n(rotation)=6, p=0.4848, U=23. **B** Same as in A, but for grid scores in the baseline trial. Mann-Whitney-Wilcoxon test, two-sided, statistical unit: mouse: n(stable)=6, n(rotation)=6, p=0.4848, U=23.

**Supplementary Figure S2:**
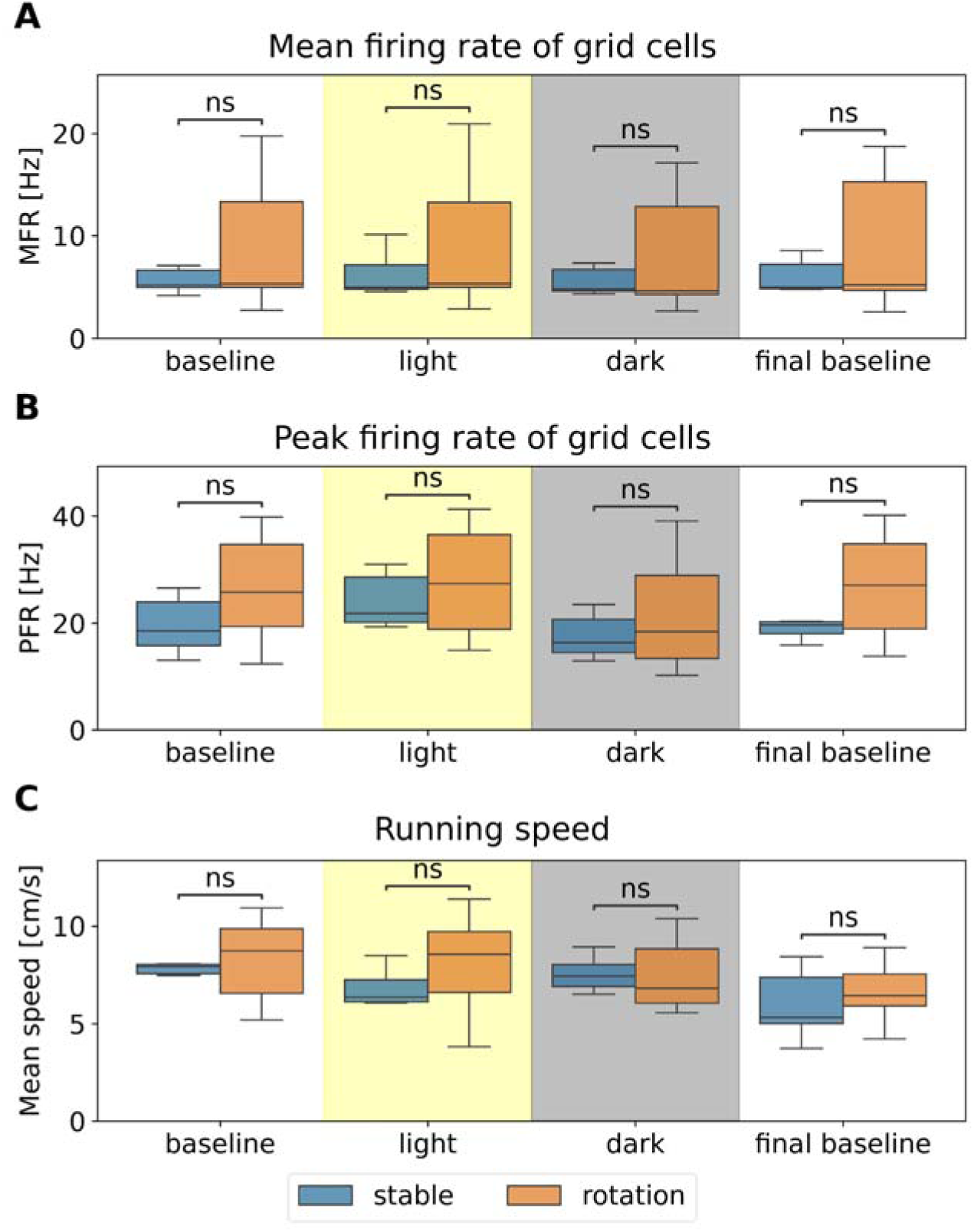
Firing activity and running speed are similar across groups and conditions. **A** Mean firing rate (MFR) of grid cells for the baseline trial, the concatenated light intervals, the concatenated dark intervals and the final baseline for the stable and the rotation group during the habituation sessions. The legend below panel C applies. Mann-Whitney-Wilcoxon test, two-sided, statistical unit: mouse; n(stable)=6, n(rotation)=6; baseline: p=0.6991, U=15; light: p=0.4848, U=13; dark: p=0.9372, U=19: final baseline: p=0.9372, U=19. **B** Peak firing rate (PFR) of grid cells for the baseline trial, the concatenated light intervals, the concatenated dark intervals and the final baseline for the stable and the rotation group during the habituation sessions. The legend below panel C applies. Mann-Whitney-Wilcoxon test, two-sided, statistical unit: mouse; n(stable)=6, n(rotation)=6; baseline: p=0.3939, U=12; light: p=0.8182, U=16; dark: p=0.8182, U=16; final baseline: p=0.4848, U=13. **C** Same for running speed. Mann-Whitney-Wilcoxon test, two-sided, statistical unit: mouse; n(stable)=6, n(rotation)=6; baseline: p=0.6991, U=15; light: p=0.3939, U=12; dark: p=0.5887, U=22: final baseline: p=0.4848, U=13.

**Supplementary Figure S3:**
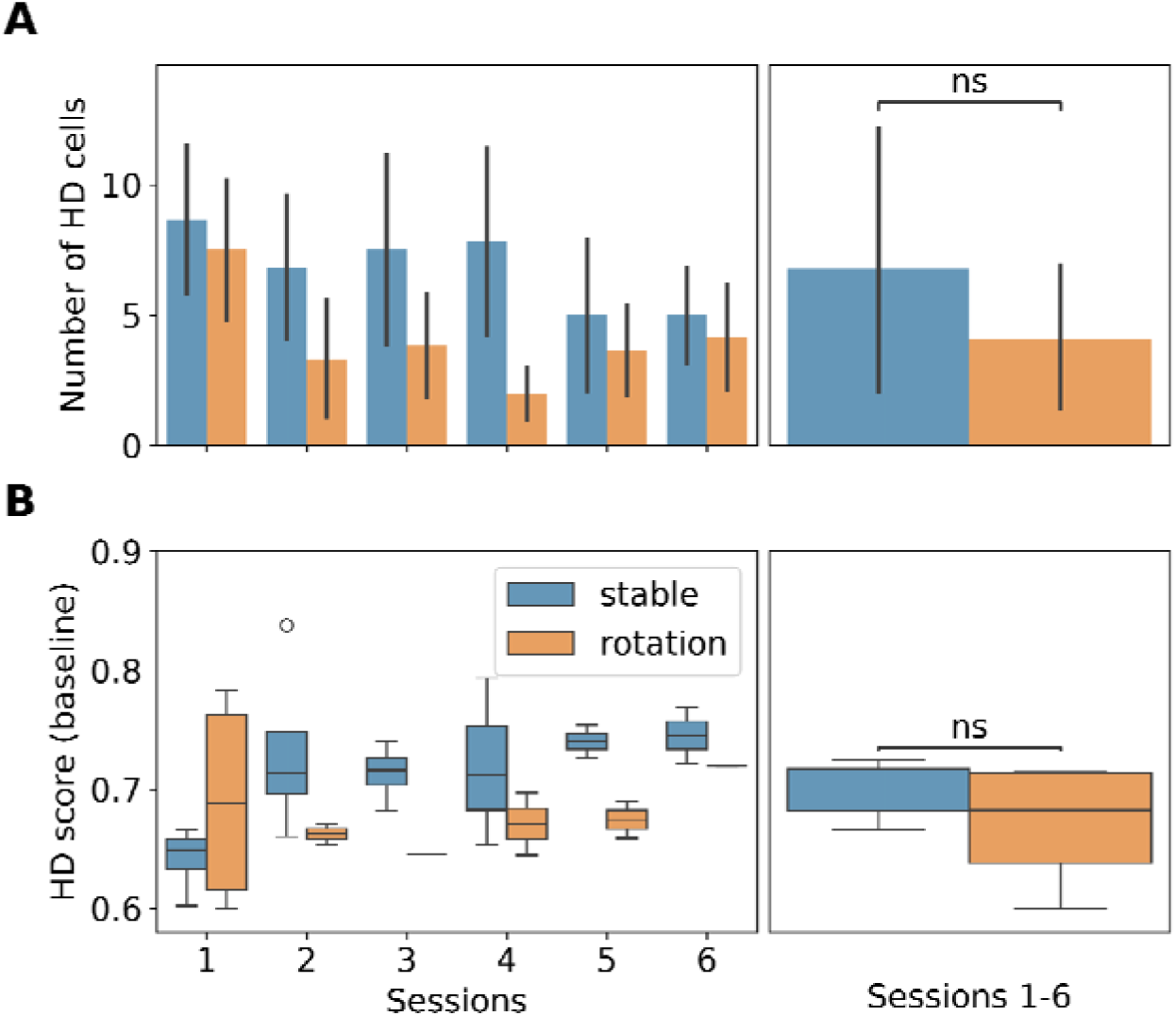
HD cell numbers and HD scores are similar across groups. **A** Left panel: Number of HD cells in each of the 6 habituation sessions for the stable and the rotation group. Right panel: Mean number of HD cells across the 6 habituation sessions. The legend from panel B applies. Mann-Whitney-Wilcoxon test, two-sided, statistical unit: mouse: n(stable)=5, n(rotation)=4, p=0.5745, U=22. **B** Same as in A, but for HD scores of HD cells in the baseline trial. Mann-Whitney-Wilcoxon test, two-sided, statistical unit: mouse: n(stable)=5, n(rotation)=4, p=0.1905, U=16.

## Acknowledgements

We thank the animal facility of the University Heidelberg for their technical assistance. We thank A. Caputi for the insightful discussions and inspiring feedback on the manuscript. This work was made possible by the following Deutsche Forschungsgemeinschaft Research Grants: the AL 1730/3-1 to KA, the CRC/TRR 1436 to HM, the CRC/TRR 384 IN-CODE to HM, the Advanced ERC grant Project-ID 101142587 to HM. We thank the Lautenschläger Foundation and the Hector Foundation for supporting the Department of Clinical Neurobiology (HM).

## Author contributions

Conceptualization: KA,HM

Methodology and experiments: PK,BT

Data analysis: PK,BT

Funding acquisition: HM,KA

Writing –original draft: BT,PK

Writing –review and editing: BT and HM with the help of the other authors

## Competing interests

Authors declare that they have no competing interests.

## Data and materials availability

The spike trains, animal position data, and task event logs will be published on the open-access Dryad platform. The computer code for data analysis will be accessible on a public GitHub repository.

## Notes

### Competing Interest Statement

The authors have declared no competing interest.

## References

1. Mittelstaedt, M.-L., and Mittelstaedt, H. (1980). Homing by Path integration in a Mammal. Naturwissenschaften.

2. Etienne, A.S., and Jeffery, K.J. (2004). Path integration in mammals. Hippocampus 14, 180–192.

3. Stangl, M., Kanitscheider, I., Riemer, M., Fiete, I., and Wolbers, T. (2020). Sources of path integration error in young and aging humans. Nat Commun 11, 2626.

4. McNaughton, B.L., Battaglia, F.P., Jensen, O., Moser, E.I., and Moser, M.-B. (2006). Path integration and the neural basis of the “cognitive map.” Nat Rev Neurosci 7, 663– 678.

5. Zhao, M., and Warren, W.H. (2015). Environmental stability modulates the role of path integration in human navigation. Cognition 142, 96–109.

6. Chen, X., McNamara, T.P., Kelly, J.W., and Wolbers, T. (2017). Cue combination in human spatial navigation. Cogn Psychol 95, 105–144.

7. Basnak, M.A., Kutschireiter, A., Okubo, T.S., Chen, A., Gorelik, P., Drugowitsch, J., and Wilson, R.I. (2025). Multimodal cue integration and learning in a neural representation of head direction. Nat Neurosci 28, 1729–1740.

8. Müller, M., and Wehner, R. (2007). Wind and sky as compass cues in desert ant navigation. Naturwissenschaften 94, 589–594.

9. Shaverdian, S., Dirlik, E., Mitchell, R., Tocco, C., Webb, B., and Dacke, M. (2022). Weighted cue integration for straight-line orientation. iScience 25, 105207.

10. Sutton, J.E. (2002). Multiple-landmark piloting in pigeons (Columba livia): landmark configuration as a discriminative cue. J Comp Psychol 116, 391–403.

11. Hébert, M., Bulla, J., Vivien, D., and Agin, V. (2017). Are Distal and Proximal Visual Cues Equally Important during Spatial Learning in Mice? A Pilot Study of Overshadowing in the Spatial Domain. Front Behav Neurosci 11, 109.

12. Biegler, R., and Morris, R.G. (1993). Landmark stability is a prerequisite for spatial but not discrimination learning. Nature 361, 631–633.

13. Fetsch, C.R., Pouget, A., DeAngelis, G.C., and Angelaki, D.E. (2011). Neural correlates of reliability-based cue weighting during multisensory integration. Nat Neurosci 15, 146–154.

14. Ernst, M.O., and Bülthoff, H.H. (2004). Merging the senses into a robust percept. Trends Cogn Sci 8, 162–169.

15. Pearce, J.M. (2009). The 36th Sir Frederick Bartlett lecture: an associative analysis of spatial learning. Q J Exp Psychol (Hove) 62, 1665–1684.

16. Yoganarasimha, D., Yu, X., and Knierim, J.J. (2006). Head direction cell representations maintain internal coherence during conflicting proximal and distal cue rotations: comparison with hippocampal place cells. J Neurosci 26, 622–631.

17. Yoder, R.M., Clark, B.J., Brown, J.E., Lamia, M.V., Valerio, S., Shinder, M.E., and Taube, J.S. (2011). Both visual and idiothetic cues contribute to head direction cell stability during navigation along complex routes. J Neurophysiol 105, 2989–3001.

18. Evans, T., Bicanski, A., Bush, D., and Burgess, N. (2016). How environment and self-motion combine in neural representations of space. J Physiol 594, 6535–6546.

19. Skaggs, W.E., Knierim, J.J., Kudrimoti, H.S., and McNaughton, B.L. (1995). A model of the neural basis of the rat’s sense of direction. Adv Neural Inf Process Syst 7, 173–180.

20. Zhang, K. (1996). Representation of spatial orientation by the intrinsic dynamics of the head-direction cell ensemble: a theory. J Neurosci 16, 2112–2126.

21. Kim, S.S., Rouault, H., Druckmann, S., and Jayaraman, V. (2017). Ring attractor dynamics in the central brain. Science 356, 849–853.

22. Chaudhuri, R., Gerçek, B., Pandey, B., Peyrache, A., and Fiete, I. (2019). The intrinsic attractor manifold and population dynamics of a canonical cognitive circuit across waking and sleep. Nat Neurosci 22, 1512–1520.

23. Shapiro, M.L., Tanila, H., and Eichenbaum, H. (1997). Cues that hippocampal place cells encode: dynamic and hierarchical representation of local and distal stimuli. Hippocampus 7, 624–642.

24. Hafting, T., Fyhn, M., Molden, S., Moser, M.-B., and Moser, E.I. (2005). Microstructure of a spatial map in the entorhinal cortex. Nature 436, 801–806.

25. Gardner, R.J., Hermansen, E., Pachitariu, M., Burak, Y., Baas, N.A., Dunn, B.A., Moser, M.-B., and Moser, E.I. (2022). Toroidal topology of population activity in grid cells. Nature 602, 123–128.

26. Peng, J.-J., Throm, B., Najafian Jazi, M., Yen, T.-Y., Pizzarelli, R., Monyer, H., and Allen, K. (2025). Grid cells accurately track movement during path integration-based navigation despite switching reference frames. Nat Neurosci 28, 2092–2105.

27. Barry, C., Bush, D., O’Keefe, J., and Burgess, N. (2012). Models of grid cells and theta oscillations. Nature 488, E1–E2; discussion E2–E3.

28. Bush, D., Barry, C., Manson, D., and Burgess, N. (2015). Using Grid Cells for Navigation. Neuron 87, 507–520.

29. Campbell, M.G., Ocko, S.A., Mallory, C.S., Low, I.I.C., Ganguli, S., and Giocomo, L.M. (2018). Principles governing the integration of landmark and self-motion cues in entorhinal cortical codes for navigation. Nat Neurosci 21, 1096–1106.

30. Barry, C., Hayman, R., Burgess, N., and Jeffery, K.J. (2007). Experience-dependent rescaling of entorhinal grids. Nat Neurosci 10, 682–684.

31. Rueckemann, J.W., Sosa, M., Giocomo, L.M., and Buffalo, E.A. (2021). The grid code for ordered experience. Nat Rev Neurosci 22, 637–649.

32. Jeffery, K.J. (1998). Learning of landmark stability and instability by hippocampal place cells. Neuropharmacology 37, 677–687.

33. Knierim, J.J., Kudrimoti, H.S., and McNaughton, B.L. (1995). Place cells, head direction cells, and the learning of landmark stability. J Neurosci 15, 1648–1659.

34. Boyle, S.C., Kayser, S.J., and Kayser, C. (2017). Neural correlates of multisensory reliability and perceptual weights emerge at early latencies during audio-visual integration. Eur J Neurosci 46, 2565–2577.

35. Jeffery, K.J., Page, H.J.I., and Stringer, S.M. (2016). Optimal cue combination and landmark-stability learning in the head direction system. J Physiol 594, 6527–6534.

36. Fetsch, C.R., DeAngelis, G.C., and Angelaki, D.E. (2013). Bridging the gap between theories of sensory cue integration and the physiology of multisensory neurons. Nat Rev Neurosci 14, 429–442.

37. Shaikh, D. (2022). Learning multisensory cue integration: A computational model of crossmodal synaptic plasticity enables reliability-based cue weighting by capturing stimulus statistics. Front Neural Circuits 16, 921453.

38. Ajabi, Z., Keinath, A.T., Wei, X.-X., and Brandon, M.P. (2023). Population dynamics of head-direction neurons during drift and reorientation. Nature 615, 892–899.

39. Stahl, J.S. (2004). Using eye movements to assess brain function in mice. Vision Res 44, 3401–3410.

40. Pérez-Escobar, J.A., Kornienko, O., Latuske, P., Kohler, L., and Allen, K. (2016). Visual landmarks sharpen grid cell metric and confer context specificity to neurons of the medial entorhinal cortex. Elife 5. 10.7554/eLife.16937.

41. Allen, K., Gil, M., Resnik, E., Toader, O., Seeburg, P., and Monyer, H. (2014). Impaired path integration and grid cell spatial periodicity in mice lacking GluA1-containing AMPA receptors. J Neurosci 34, 6245–6259.

42. Dannenberg, H., Lazaro, H., Nambiar, P., Hoyland, A., and Hasselmo, M.E. (2020). Effects of visual inputs on neural dynamics for coding of location and running speed in medial entorhinal cortex. Elife 9. 10.7554/eLife.62500.

43. Chen, G., Manson, D., Cacucci, F., and Wills, T.J. (2016). Absence of Visual Input Results in the Disruption of Grid Cell Firing in the Mouse. Curr Biol 26, 2335– 2342.

44. Waaga, T., Agmon, H., Normand, V.A., Nagelhus, A., Gardner, R.J., Moser, M.-B., Moser, E.I., and Burak, Y. (2022). Grid-cell modules remain coordinated when neural activity is dissociated from external sensory cues. Neuron 110, 1843–1856.e6.

45. Sargolini, F., Fyhn, M., Hafting, T., McNaughton, B.L., Witter, M.P., Moser, M.-B., and Moser, E.I. (2006). Conjunctive representation of position, direction, and velocity in entorhinal cortex. Science 312, 758–762.

46. Jeffery, K.J., and O’Keefe, J.M. (1999). Learned interaction of visual and idiothetic cues in the control of place field orientation. Exp Brain Res 127, 151–161.

47. Chakraborty, S., Anderson, M.I., Chaudhry, A.M., Mumford, J.C., and Jeffery, K.J. (2004). Context-independent directional cue learning by hippocampal place cells. Eur J Neurosci 20, 281–292.

